# Oscillatory Hedgehog signaling and temporal coordination of *atonal* dynamics at the eye differentiation front in *Drosophila*

**DOI:** 10.64898/2026.03.16.712009

**Authors:** Minh-Son Phan, Claire Mestdagh, Francois Schweisguth

## Abstract

Pattern formation in the *Drosophila* eye involves the pulsatile expression of the proneural gene *atonal* (*ato*) and the periodic activation of Notch. Oscillatory expression of the *ato* gene results from the synchronous activation of two distinct *ato* enhancers that are active in two adjacent rows of cell clusters along the front of differentiation. How synchronous enhancer activation is achieved along the front is not known. Hedgehog (Hh) is a diffusive signal produced just posterior to the front which contributes to regulate *ato* gene expression. Here, we find that both *ato* enhancers are regulated by Hh and that two Hh target genes, *patched* (*ptc*) and *decapentaplegic* (*dpp*), are expressed at the front of differentiation in an oscillatory manner downstream of a constant Hh signal. Consistent with a role of Hh in the temporal coordination of *ato* gene expression along the front, lowering Hh activity was associated with pattern irregularities along the front. We propose a model whereby the periodic expression of the Hh receptor Ptc under the control of pulsatile Ato and/or Notch dynamics produces rhythmic changes in extracellular Hh which would in turn coordinate *ato* dynamics locally. In this model, oscillatory Hh would provide a temporal cue to coordinate patterning dynamics along the front of differentiation in the developing eye.

## Introduction

The formation of reproducible patterns of cell fates relies on the interpretation of positional cues as well as on local cell-cell interactions mediating self-organization (Green and Sharpe, 2015; Schweisguth and Corson, 2019). Self-organization based on inhibitory cell-cell interactions by Delta-Notch signaling generates disordered arrays of cell fates (Collier et al., 1996; Phan et al., 2024; Sánchez-Iranzo et al., 2022; Simpson, 1990). When guided by tissue geometry and/or positional cues, self-organized Notch dynamics can also produce more elaborate patterns (Cohen et al., 2020; Corson et al., 2017). In the developing eye of *Drosophila*, Notch-based self-organization produces a crystal-like array of R8 photoreceptor cells (Couturier et al., 2026; Roignant and Treisman, 2009; Warren and Kumar, 2023). In this neuroepithelium, cell fate patterning takes place at a front of differentiation that sweeps through the eye primordium, with R8 cells emerging in a locally synchronous manner in the wake of the moving front to produce regular rows of R8 cells (Couturier et al., 2026; Roignant and Treisman, 2009; Warren and Kumar, 2023). The quasi-synchronous emergence of the R8 cells results from the local synchrony in Notch dynamics along the front of differentiation (Couturier et al., 2026). How Notch dynamics is temporally coordinated at the tissue level to ensure local synchrony is not known. In many tissues, the timing of cell differentiation is coordinated at the tissue scale via extrinsic timing cues. However, the repetitive nature of the emergence of sequential rows of R8 cells make this model unlikely, and temporal cues are more likely to be embedded within the patterning mechanism itself.

The formation of the R8 pattern in the developing fly eye has been extensively studied (Roignant and Treisman, 2009; Warren and Kumar, 2023). The adult eye comprises ∼750 multicellular light-receiving units, or ommatidia, and each of these ommatidia stems from a single founder R8 cell that is singled out by Notch-mediated lateral inhibition from a group of proneural cells expressing the transcription factor Atonal (Ato), the key eye-determining gene in *Drosophila* (Baker et al., 1996; Dokucu et al., 1996; Jarman et al., 1994). These proneural clusters, known as Intermediate Groups (IGs), appear at the level of the moving front of differentiation and resolve into rows of R8s (Fig 1A). Recently, live imaging revealed oscillatory *ato* gene expression at the front of differentiation, with a new pulse of Ato expression observed each time a new row of IGs emerge while R8s become selected from within IGs during the interpulse (Couturier et al., 2026). These pulses of *ato* gene expression were shown to be produced by two distinct cis-regulatory enhancers. A first *ato5’* enhancer directs the pulse of Ato in IGs whereas a second *ato3’* enhancer regulates *ato* expression in groups of cells located anterior to the IG row and known as the Initial Clusters (ICs) (Couturier et al., 2026; Sun et al., 1998)(Fig 1A,B). Thus, these two *ato* enhancers are active at the same time but in different cells to produce periodic Ato expression at the tissue scale (Couturier et al., 2026). How temporal coordination is achieved to ensure that the activation of the *ato5’* and *ato3’* enhancers is locally synchronous along the front, thereby ensuring regular patterning dynamics at the tissue level, is not known.

**Figure 1.**
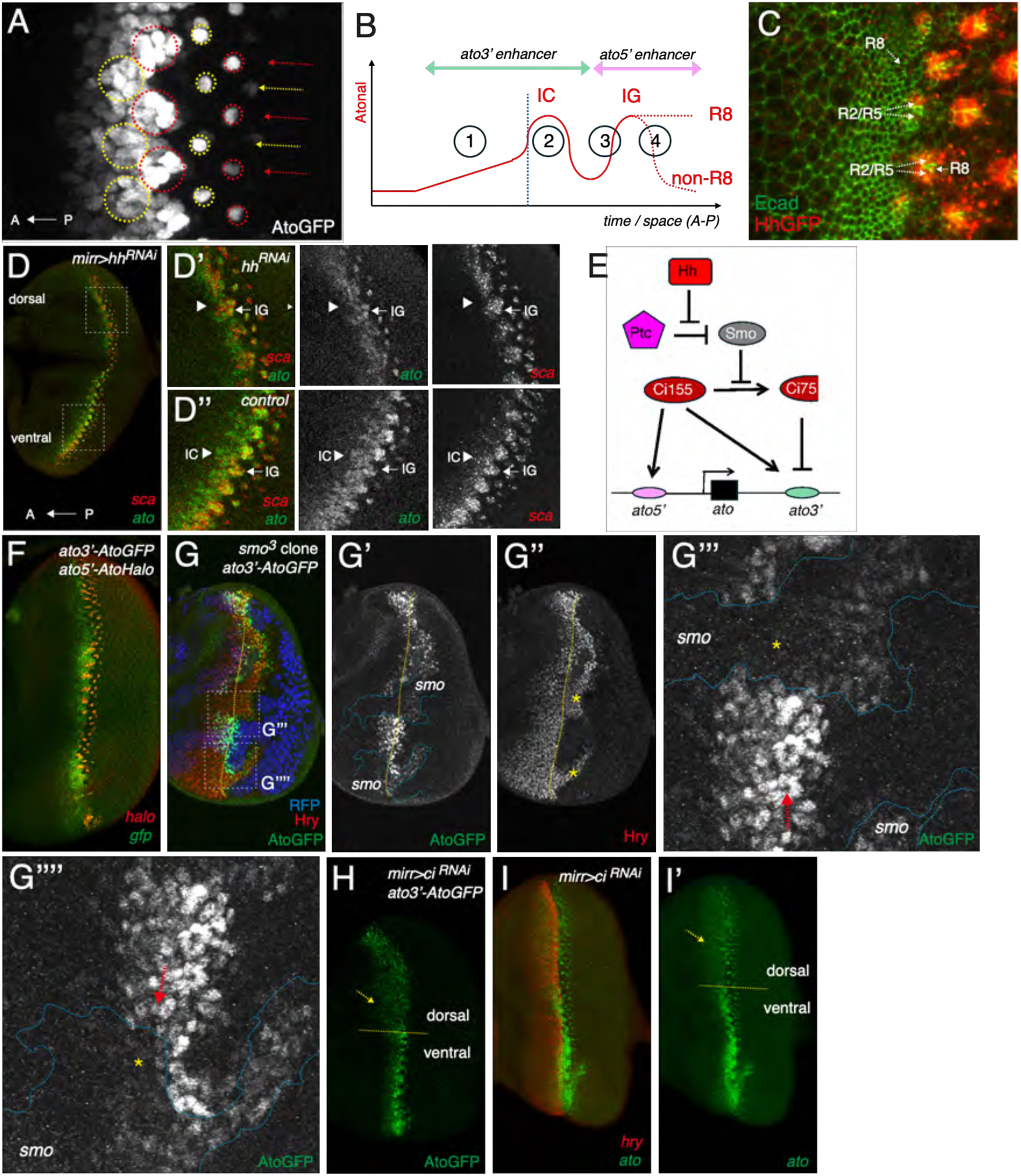
Hh regulates *ato* gene expression in IGs and ICs. A) Pattern of Ato accumulation (AtoGFP, white) at the MF. Clusters of Ato-expressing cells define rows of ICs (large yellow circles) and IGs (large red circles). These clusters are organized into columns along the A-P axis (arrows); ICs, IGs and R8s (small circles) of alternate columns are color-coded (in yellow and red). Low Ato levels were detected anterior to the ICs. In this and all other images, anterior is left and dorsal is top. B) Schematics showing the temporal profile of Ato accumulation. The *ato3’* enhancer regulates the initiation of *ato* gene expression (1) and its up-regulation in ICs (2). The *ato5’* enhancer regulates *ato* gene expression in IGs (3) and mediates its restriction to R8s (4). The vertical blue line marks the down-regulation of Hairy (Hry) and the up-regulation of the E(spl)-HLH factors, marking the onset of proneural differentiation. C) GFP-tagged Hh (anti-GFP, red; E-Cadherin, noted Ecad, green) was first detected in the R2/R5 cells that flank the R8s along the posterior edge of the MF. R8, R2 and R5 cells were identified based on their position and morphology. D-D’’) Lowering Hh in the dorsal compartment of *mirr>hh^RNAi^* discs led to reduced *ato* gene expression (*ato* mRNAs, green) in ICs and IGs (high magnification panels in D). The *scabrous* (*sca*) transcripts (red) were used to detect IGs). E) Schematics of the Hh signaling pathway. The repression of Smo by Ptc is relieved upon the binding of Hh to Ptc. Smo signals intracellularly to prevent the partial proteolysis of Ci^Act^ (Ci155) into Ci^Rep^ (Ci75). Ci^Rep^ prevents the early activation of the *ato3’* enhancer whereas Ci^Act^ up-regulates the activities of the *ato3’* and *ato5’* enhancers in ICs and IGs, respectively. F) Pattern of *ato3’* and *ato5’* enhancer activities. The *ato3’* enhancer is active anterior to the MF and in ICs (*gfp* mRNAs, green) whereas the *ato5’* enhancer is active in IGs and R8 cells (*halo* mRNAs, red) (Couturier et al., 2026). G-G’’’’) Positive regulation of *ato3’* by Smo. The activation of the *ato3’* enhancer (AtoGFP, green in G) was strongly delayed in *smo* mutant clones (G-G’’’’: mutant cells (marked by the loss of nuclear RFP, blue in G; clone boundaries indicated by blue dotted lines) expressed low levels of AtoGFP posterior to the predicted position of the MF (yellow dotted line in G-G’’).The *ato3’* enhancer was not activated in *smo* mutant cells (stars in G’’’,G’’’’; ICs were indicated by red arrows in G’’’,G’’’’), indicating that Smo is required for the up-regulation of *ato3’* in ICs. Also, Hry (red, G) was not down-regulated in the absence of Smo (star, G’’). H-I’) Loss of Ci in the dorsal compartment of *mirr>ci^RNAi^* discs led to the early and ectopic activation of the *ato3’* enhancer (AtoGFP, green in H) and expression of *ato* mRNAs (green in I,I’; *hry* mRNAs, red) in cells anterior to the MF. The *ato3’* enhancer was not up-regulated in ICs (H) and the expression of *ato* remained weak in *ci^RNAi^* ICs/IGs (I; a yellow line indicates the dorsal-ventral boundary in H,I’).

The regulation of *ato* gene expression in the eye imaginal disc has been well studied. Its expression is initiated anterior to the front in a Notch-independent manner (Couturier et al., 2026; Sun et al., 1998). Then, in response to a Delta signal produced by the late IG cells of the same column, it is up-regulated in ICs (Couturier et al., 2026; Sun et al., 1998). The activity of the *ato3’* enhancer is up-regulated in ICs by nuclear activated Notch via the sequence-specific DNA-binding factor Suppressor of Hairless (Couturier et al., 2026; Henrique and Schweisguth, 2019; Kopan and Ilagan, 2009). Since Delta appears to be transiently produced by late IG cells, this signal only produces a pulse of Notch activity in ICs, hence a pulse of Ato (Couturier et al., 2026). Ato is then re-expressed at high levels by the same cells that now form the IGs, this time under the control of the *ato5’* enhancer, to contribute to the next Ato pulse. This regulatory logic repeats itself as IGs resolve via Notch-mediated lateral inhibition, with late IG cells producing a transient Delta signal, up-regulating Ato expression in anterior cells which will form the ICs. Thus, in this relay model, patterning cues self-propagate in a column-autonomous manner, with Delta produced in late IGs positively regulating *ato* gene expression in ICs (Couturier et al., 2026). Although this self-propagating mechanism appeared to operate within columns of cells, the observation of Ato oscillations at the tissue scale strongly suggests that a coordination mechanism acts to ensure the synchrony in *ato* gene expression in neighboring columns, thereby promoting the smooth progression of the differentiation front.

Here, we addressed whether signaling by Hedgehog (Hh) contributes to coordinating the activities of the *ato3’* and *ato5’* enhancers to produce Ato oscillations at the tissue scale. We showed that these two enhancers are regulated by Hh signaling and found that two Hh target genes, *patched* (*ptc*) and *decapentaplegic* (*dpp*), are expressed in an oscillatory manner at the front of differentiation downstream of a constant Hh signal. Since lowering Hh activity led to rare and mild patterning defects, we propose that Hh contributes to coordinate *ato* dynamics across columns.

## Results

### Hh regulates *ato* gene expression in ICs and IGs

Hh is expressed posterior to the MF by the R2 and R5 cells (Ma et al., 1993) and diffuses to regulate gene expression in cells located within and anterior to the MF. Hh is known to derepress the expression of the *dpp* and *ato* genes in these cells, notably by antagonizing the production of the repressive form of Cubitus interruptus (Ci) known as Ci^Rep^ (or Ci75) (Borod and Heberlein, 1998; Domínguez and Hafen, 1997; Fu and Baker, 2003; Pappu et al., 2003; Strutt and Mlodzik, 1997). Here, we confirmed these findings and showed that a functional BAC-encoded HhGFP, used here to detect Hh, was first detected posterior to the emerging R8 cells within the same column (Fig 1C) and that the silencing of the *hh* gene in the dorsal compartment, using a *mirror-Gal4* (*mirr-Gal4*) driver, led to reduced *ato* expression in both ICs and IGs (Fig 1D-D’’). Consistent with earlier findings (Fu and Baker, 2003; Greenwood and Struhl, 1999), a complete loss of Hh signaling in cells mutant for the Hh-signal transduction protein Smoothened (Smo) largely stopped the progression of the differentiation front, as seen by the expression of Hairy (Hry) in mutant cells (Fig 1G-G’’). Using an *ato3’-AtoGFP* transgene to study the initiation of *ato* gene expression and its up-regulation in ICs (Fig 1F) (Couturier et al., 2026), we found that signaling by Smo is required for the activity of the *ato3’* enhancer in the MF (Fig 1G’’’’). Of note, the loss of *ato3’* activity seen in *smo* mutant clones could either result from an increase in Ci^Rep^ level or from a loss of the activator form of Ci, Ci^Act^ (or Ci155), which is produced in the presence of Hh (Fig 1E). To distinguish between these two possibilities, we examined the effect of the silencing of the *ci* gene on *ato3’* enhancer activity. Early and ectopic *ato3’* enhancer activity and *ato* gene expression were observed in cells anterior to the MF in *mirr>ci^RNAi^* discs whereas no up-regulation of *ato* gene expression was observed in the MF (Fig 1H-I’). This suggested that Ci^Rep^ acts via the *ato3’* enhancer to delay the onset of *ato* expression and that Ci^Act^ is required to up-regulate *ato* gene expression in ICs and IGs. Since Ci binding sites were predicted at the *ato* locus (Gurdziel et al., 2015) and since both Ci^Rep^ and Ci^Act^ were found to physically interact with cis-regulatory sequences 5’ and 3’ to the *ato* gene (Biehs et al., 2010), we propose that Hh regulates *ato* gene expression in both ICs and IGs via Ci. Additionally, Hh might also regulate *ato* gene expression indirectly via the transcription factors of the Retina Genetic Network (Brás-Pereira et al., 2016; Pappu et al., 2003; Zhang et al., 2006b). Thus, Hh regulates *ato* gene expression in both ICs and IGs.

### Reconstructing temporal dynamics of gene expression from fixed samples

Since Hh is a diffusive signal, it might coordinate the concomittant activation of the *ato3’* and *ato5’* enhancers in distinct ICs and IGs along the front. A simple hypothesis would be that a pulsatile Hh signal acts as a temporal coordination cue. To test whether the Hh signal may be pulsatile, we first examined the transcription dynamics of well-characterized Hh target genes. However, studying gene expression on fixed samples has inherent limitations to infer temporal dynamics. To circumvent these limitations, we used the shift in synchrony revealed by the traveling waves of Ato (Fig 2A) to reconstruct the expression dynamics of Hh target genes using single molecule inexpensive fluorescent in situ hybridization (smiFISH) (Couturier et al., 2026). As a proof of principle, we first reconstructed the periodic spatial-temporal expression of the *ato* and *E(spl)m*δ*-HLH* (*m*δ) genes from fixed samples in discs expressing GFP-tagged mδ (GFPmδ) (Couturier et al., 2026). Following maximum z-projection of the *ato* smiFISH and anti-GFP signals, we manually annotated the positions of the Ato clusters along the MF and categorized them in 8 stages covering two periods of the Ato cycle, from an early IC stage to an R8 stage (Fig 2A’,A’’ and Fig S1). The dynamics of Ato at intervening columns was obtained by interpolating positions and annotating categories automatically by introducing a one period shift (Fig S1). Annotated clusters were then oriented perpendicular to the MF, cropped and intensity normalized (Fig S1). This pipeline produced a discretized average dynamic of Ato and E(spl)mδ over two consecutives pulses of Ato (Fig 2B). We next realigned these 8 frames along the AP axis to compensate for the progression of the MF, assigned a time duration for each of the 8 categories based on their relative frequency and smoothened the signals in time to reduce variability (Fig S1). This approach established the average spatial-temporal dynamics of Ato and E(spl)mδ at the MF (movie 1).

**Figure 2.**
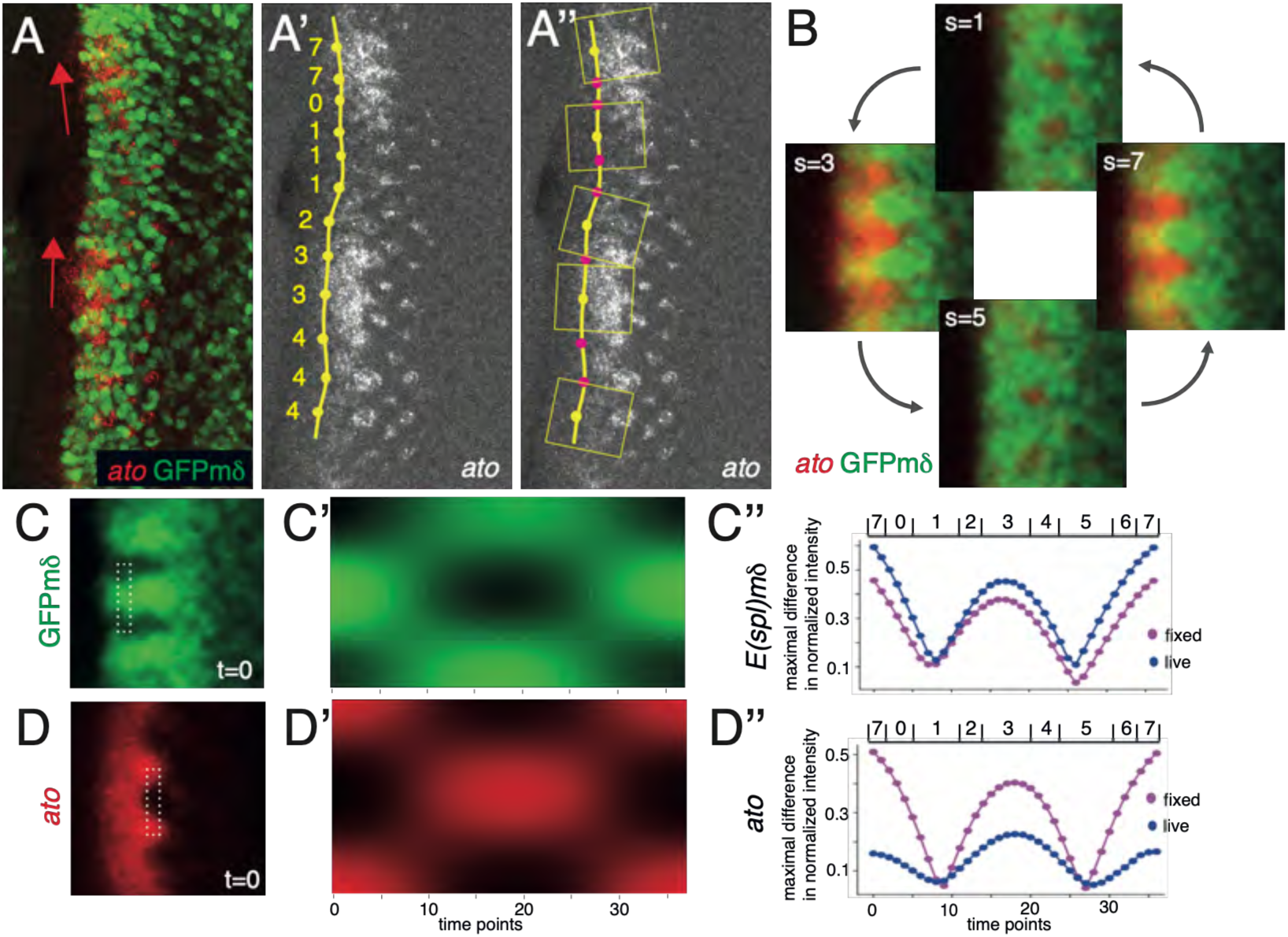
From snapshots to dynamic gene expression. A-A’’) Accumulation patterns of *ato* mRNA (red in A) and GFPmδ (anti-GFP, green in A). Two consecutive pulses of Ato are observed (arrows in A). The Ato clusters were manually annotated and staged (A’; see Methods), then oriented perpendicular to the front and cropped (represented as yellow boxes in A’’; only one every third clusters are shown for better visualization). B) Averaged views of the Ato cluster at four different stages (out of 8), spanning two Ato consecutive oscillations C-D’’) The Ato and E(spl) dynamics that were reconstructed from fixed samples reproduced the average dynamics measured by live imaging. Kymographs (C’,D’) were produced from GFPmδ and *ato* pseudo-time series (C,D; boxed area indicate the ROIs used to generate kymographs; see Methods). The expression dynamics obtained directly from live imaging (Couturier et al., 2026) (GFPmδ in C’’, and AtoGFP in D’’) was compared to those obtained from fixed samples (GFPmδ in C’’, *ato*, D’’) by plotting over time the maximal difference measured between all normalized values at each time point of the corresponding kymographs (C’,D’: kymographs produced with data from fixed samples).

Kymograph analysis showed that these average dynamics were very similar to the average dynamics obtained by live imaging AtoGFP and GFPmδ in *ex vivo* cultured eye discs (Couturier et al., 2026) (Fig 2C-D’’). We conclude that gene expression dynamics at the MF could be faithfully reconstructed from fixed samples and that the expression patterns of *ato* and GFPmδ provided enough information to reconstruct patterning dynamics at the MF. We next used this approach to study the transcription dynamics of several Hh target genes.

### Oscillatory expression of *ptc* and *dpp* at the MF

To study the transcription dynamics of the *<u>ptc</u>* gene, a direct target of Hh (Alexandre et al., 1996; Parker et al., 2011), we used a *ptc* intronic probe that detected nuclear nascent transcripts as bright foci revealing the synthesis of primary mRNAs at the gene locus. Since intronic probes detect sites of RNA synthesis, the intensity of the fluorescence signal can serve to measure the level of primary transcription. Using the *ptc* smiFISH signal as a readout of transcription in fixed eye discs, we observed that the transcription of the *ptc* is dynamic at the MF, with increased transcription of *ptc* matching the pulses of *ato* expression (Fig 3A,A’). Using the *ato* and GFPmδ signals to register the clusters in time and space, we reconstructed the average dynamics of *ptc* (Fig 3B), we found that the expression of *ptc* appeared to oscillate in MF cells (Fig 3B’, movies 2,3). Kymograph analysis further showed that the transcriptional oscillations of the *ptc* and *ato* genes had the same period and phase. This indicated that the *ptc* gene is expressed in an oscillatory manner at the MF.

**Figure 3.**
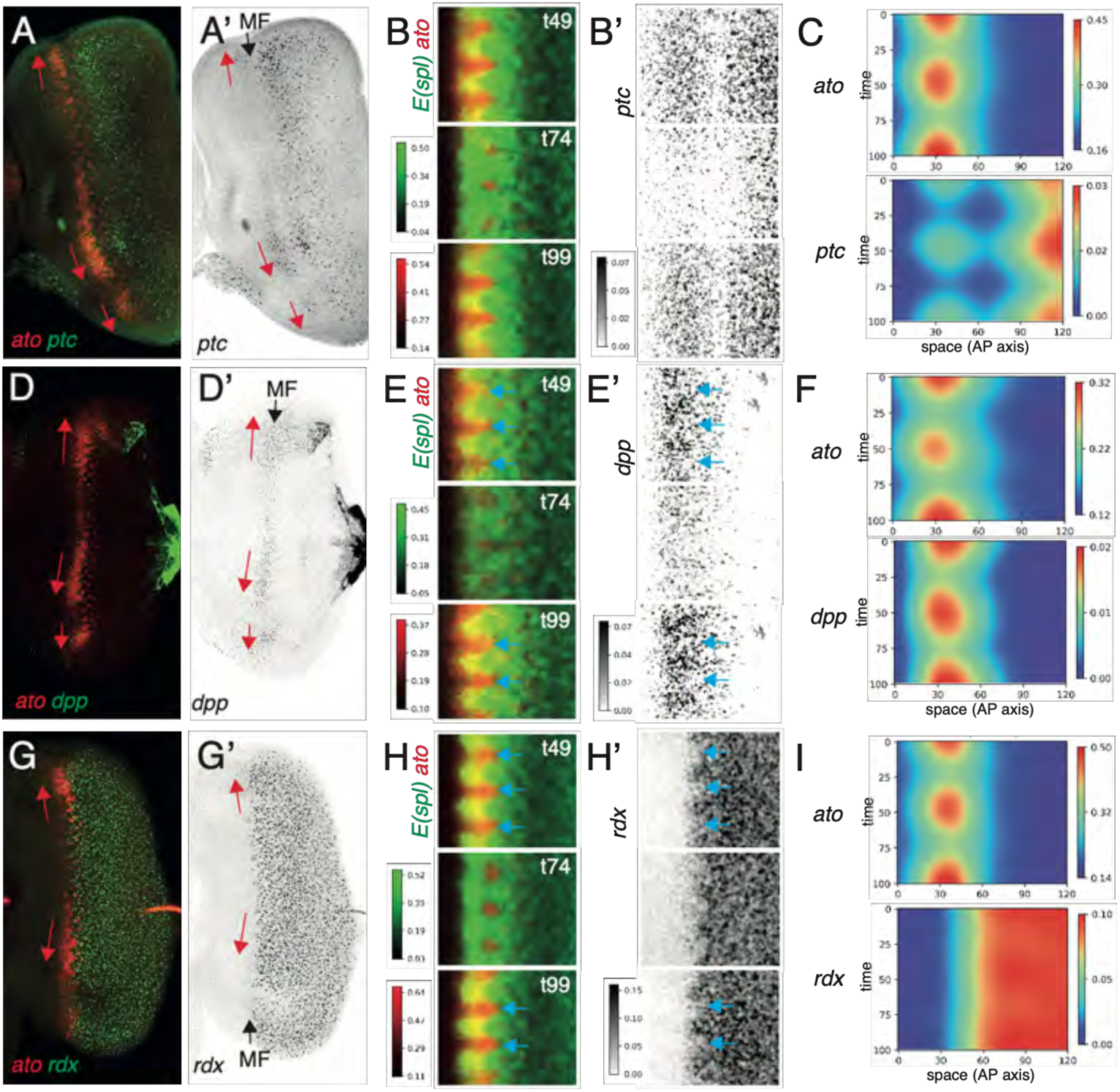
Oscillatory expression of the *ptc* and *dpp* genes. A,A’) The transcription patterns of the *ptc* (green) and *ato* genes (red) were studied by smiFISH in GFPmδ/+ discs (the GFPmδ signal, not shown here, was used together with *ato* for reconstructing the dynamics of *ptc* transcription). Increased transcription of *ptc* was seen in MF cells expressing high *ato* levels. B,B’) Dynamics of *ptc* gene transcription (B’) based on the dynamics of *ato* and *E(spl)* expression reconstructed from fixed samples (B) (n=144 clusters, from n=9 discs). C) Kymographs showing the synchronous pulsatile expression of *ato* and *ptc* expression at the MF (time in arbitrary units, space in pixels). D-I) Dynamics of *dpp* (D-F; n=88 clusters, from n=5 discs) and *rdx* gene transcription (G-I; n=172 clusters, from n=14 discs; panels as in A-C). Oscillatory expression was observed for *dpp* in the MF (D-F; note the increased *dpp* expression in IGs, see blue arrows in E,E’). In contrast, no oscillatory expression was observed for the *rdx* gene, that is expressed posterior to the MF (G-I). No *rdx* gene expression was detected in IGs (blue arrows in H,H’); the onset of *rdx* expression was concomitant to R8 selection.

To possibly extend this unexpected observation to other Hh target genes, we studied the transcription dynamics of the *dpp* and *rdx* genes. The *dpp* gene is expressed in the MF downstream of Hh (Heberlein et al., 1995; Pan and Rubin, 1995; Strutt et al., 1995), whereas the *rdx* gene is expressed posterior to the MF and is regulated by Hh and EGF (Kent et al., 2006). Here, we confirmed that the *dpp* gene is expressed at the level of the MF and observed increased level of *dpp* transcription in cells with high *ato* expression (Fig 3D,D’), suggestive of pulsatile *dpp* expression (of note, the uneven expression of *dpp* in the furrow had been noticed earlier (Domínguez, 1999)). Reconstructing *dpp* dynamics from fixed samples again showed an oscillatory pattern of *dpp* expression in synchrony with the Ato pulses (Fig 3E-F). Unlike *ptc*, however, low *dpp* transcription was observed in IGs (arrows in Fig 3E,E’; see Fig S2D,D’). Thus, the *dpp* gene is expressed in an oscillatory manner at the MF and its expression in the MF appears to be spatially regulated (movies 4,5).

We next studied the *rdx* gene and confirmed that it is expressed posterior to the MF (Zhang et al., 2006a) (Fig 3G,G’). Analysis of *rdx* transcription dynamics indicated that this gene is expressed in a relatively constant manner over time in cells posterior to the MF (Fig 3H-I).

Our analysis further showed that the *rdx* gene is transcribed only after IGs have resolved (movie 6; arrows in Fig 3H,H’). We therefore conclude that not all Hh target genes are expressed in an oscillatory manner in the eye, implying that additional regulatory inputs act to produce, or prevent, oscillations of Hh target gene expression at the MF.

### Non-pulsative Hh produced oscillatory expression of the *ptc* gene

We next asked whether the oscillatory pattern of *ptc* and *dpp* transcription resulted from the production of a periodic Hh signal. To test this hypothesis, we examined the dynamics of *hh* transcription using an intronic probe. Applying our temporal-spatial registration pipeline to fixed samples we found that the *hh* gene was expressed in a relatively constant manner in cells posterior to the front of differentiation (Fig 4A-B, movie 7). Thus, the oscillatory expression of *ptc* and *dpp* did not appear to result from an oscillatory input. This can be illustrated by registering in time and space the different dynamics using the *ato* and GFPmδ signals as templates: comparing *hh* dynamics with those of *ptc* and *dpp* clearly indicated that the oscillatory expression of the *ptc* and *dpp* genes is unlikely to result from an oscillatory Hh signal (Fig S2A,B). These observations in turn suggested that oscillatory gene expression could be generated downstream of a constant Hh signal in the eye. To test this, we examined whether a constant source of Hh could produce an oscillatory output at the MF by replacing the endogenous source of Hh by a ligand produced constitutively by differentiated cells posterior to the MF. To do so, we combined the eye-specific *hh* loss of function mutations *hh^fse^* and *hh^bar3^* (Rogers et al., 2005), thereby eliminating the endogenous source of Hh, with a GMR-Gal4 directing the expression of a UAS-Hh-GFP transgene in cells posterior to the MF, thereby rescuing the loss of *hh* activity in the eye (Chanut et al., 2000) (Fig S3). Analysis of the temporal dynamics of *ptc* transcription in this context showed oscillatory expression of the *ptc* gene (Fig 4C-D). We therefore conclude that a constant Hh signal can be associated with oscillatory *ptc* expression and propose that the oscillatory expression of *ptc* and *dpp* genes in the eye did not result from a pulsatile Hh signal.

**Figure 4.**
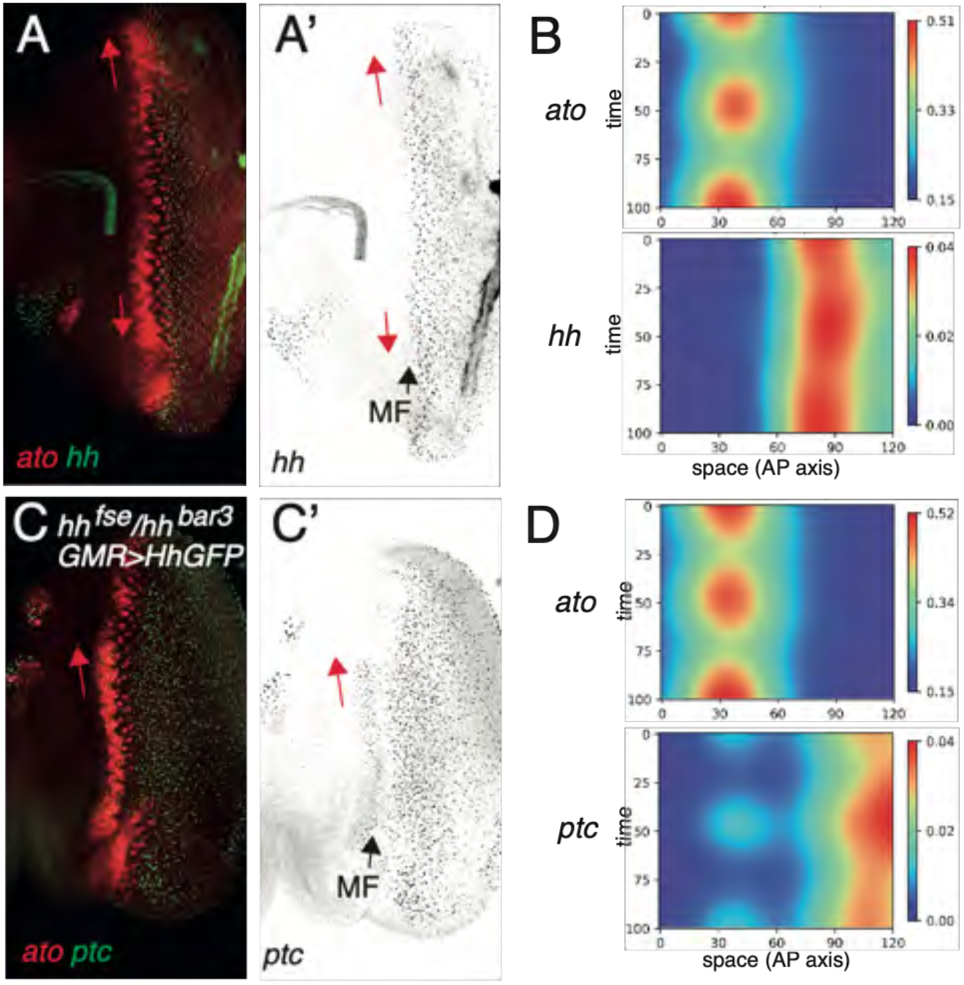
An oscillatory output produced by a non-pulsatile Hh signal. A,A’) The *hh* smiFISH signal (A,A’) detected just posterior to the MF appeared to be uniform along the MF in control discs (MF position, black arrow). In contrast, pulses of *ato* gene expression were observed (red arrows). B) Kymographs showing that *hh* transcription remained largely constant whereas pulses of *ato* transcription were measured (time in arbitrary units, space in pixels; n=268 clusters, from n=12 discs). C,C’) Expression of HhGFP by differentiated cells using *GMR>HhGFP* rescued the eye-specific loss of Hh in *hh^fse^*/*hh^bar3^* (see Fig S2) and restored the transcription of the *ptc* gene along the MF (C,C’; MF position, black arrow) with elevated *ptc* transcription observed in cells undergoing an *ato* pulse (red arrow). D) Kymographs showing the transcription dynamics of the *ato* and *ptc* genes in *hh^fse^*/*hh^bar3^ GMR>HhGFP* discs. The constant production of HhGFP by differentiated cells rescued proneural dynamics and *ptc* oscillatory gene transcription in *hh* mutant eye discs (time in arbitrary units, space in pixels; n=146 clusters, from n=9 discs).

### Reduced Ptc activity did not prevent oscillatory gene expression

Asking how a pulsatile output might be generated from a constant input, we wondered whether oscillatory gene expression is produced by a negative feedback mechanism that incorporates a time delay, as seen in other oscillatory processes that rely on a molecular clock (Harima et al., 2014; Miao and Pourquié, 2024). Indeed, the inhibition of Smo by Ptc might in principle underlie oscillatory *ptc* gene expression via a negative feedback mechanism. We note, however, that the negative regulation of Smo by Ptc is known to repress pathway activity in the absence of Hh and reduce it in the presence of Hh (Briscoe and Thérond, 2005; Ingham et al., 1991), to limit excess Hh diffusion (Chen and Struhl, 1996) and to possibly gate the response to Hh over time (Chou et al., 2010) but Ptc is not known to generate oscillations. Nevertheless, we addressed this possibility by first asking whether *ptc* transcription dynamics translated into pulses of Ptc protein accumulation. Using an anti-Ptc antibody, we detected low levels of Ptc in MF cells (Fig 5A,A’). Ptc appeared to localize into intracellular dots that likely correspond to endosomes (Martìn et al., 2001). Increased accumulation of Ptc was detected along the front at positions where Sens-positive IG cells were detected (Fig 5A). This suggested that the oscillatory expression of the *ptc* gene resulted in a periodic pattern of Ptc.

**Figure 5.**
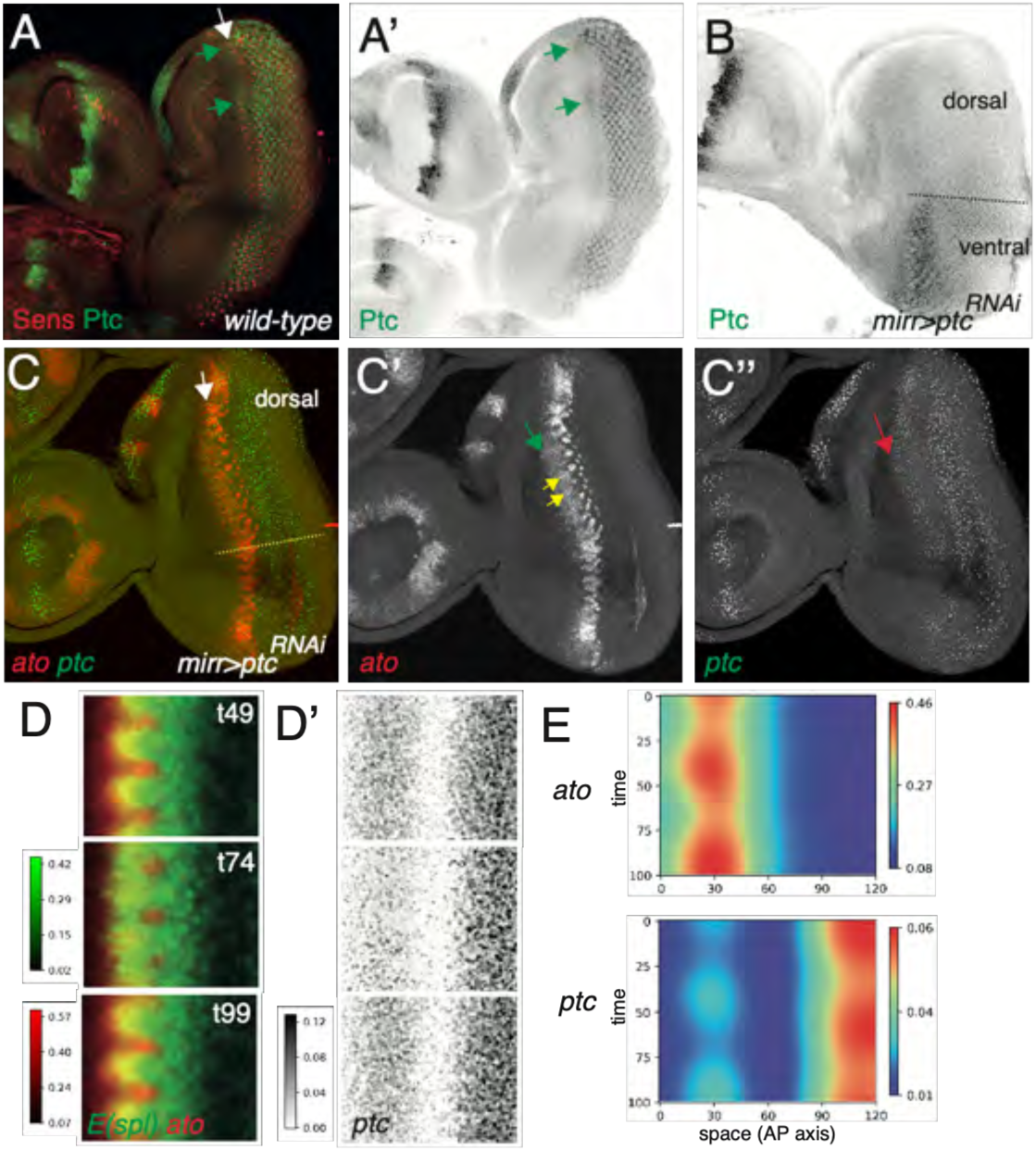
Oscillatory *ptc* transcription upon reduced Ptc activity. A-B) Pattern of Ptc protein accumulation in wild-type (A,A’) and *mirr>ptc^RNAi^* discs (B). Higher levels of Ptc were detected close to Sens-positive IG cells in control discs (green arrows in A,A’; MF indicated by a white arrow in A). The silencing of *ptc* in the dorsal compartment led to a strong decrease in Ptc protein levels in *mirr>ptc^RNAi^* discs (B). C-C’’) Patterns of *ptc* (C,C’’, green) and *ato* (C,C’, red) transcription in *mirr>ptc^RNAi^* discs (expressing also GFPmδ, not shown). Increased Hh signaling resulting from low Ptc activity led to ectopic (green arrow, C’) expression of the *ato* and *ptc* genes anterior to the MF (white arrow in C). Additionally, *ato* transcription appeared to persist in late ICs at a stage when IGs have resolved into R8s (yellow arrows, C’). Note that the silencing of the *ptc* gene did not affect the detection of *ptc* transcription foci using intronic probes (C’’). D-E) Dynamics of *ptc* gene transcription in *mirr>ptc^RNAi^* discs (D-E; n=153 clusters, from n=13 discs; data analyzed as in Fig 3). Oscillatory transcription was observed in the MF for the *ato* and *ptc* genes. Note the ectopic expression of *ato* anteriorly and the altered transcription dynamics (compare with Fig 3C). The dynamics of *ptc* and *ato* transcription appeared to be similarly altered, consistent with Ato contributing to *ptc* gene regulation at the MF.

We next tested whether the activity of Ptc was essential to generate oscillatory gene expression at the MF. The silencing of *ptc* led to a strong loss of Ptc protein (Fig 5B) and to the ectopic transcription of the *ato* and *ptc* genes in anterior cells away from the MF (Fig 5C-C’’; see green and red arrows in panels C’ and C’’, respectively), indicative of increased diffusion of Hh and/or increased Hh signaling activity. We also observed that the expression of *ato* appeared to persist in late ICs, at a stage when R8 cells emerged from late IGs (Fig 5C’). This prolonged expression of *ato* is consistent with a role of Ptc in down-regulating the Hh-dependent regulation of the *ato3’* enhancer in ICs. Thus, the activity of Ptc appeared to be strongly reduced in *mirr>ptc^RNAi^* discs [note that a complete loss of *ptc* activity in clones was shown to result in ectopic Ato expression (Domínguez, 1999)]. Despite a strong down-regulation of Ptc, kymograph analysis of the reconstructed transcription dynamics showed that the *ato* and *ptc* genes were still expressed in an oscillatory manner in *mirr>ptc^RNAi^* discs (Fig 5D-E). This observation did not support the view that a Ptc-mediated negative feedback mechanism produces oscillatory *ptc* expression.

An alternative possibility is that the oscillatory expression of the *ptc* and *dpp* genes results from their regulation by transcription factors that are active in a pulsatile manner in the MF, e.g. Ato and E(spl). Three observations supported this hypothesis. First, not all Hh target genes are expressed in an oscillatory manner in the eye (Fig 3). Second, the pulses of *dpp* expression showed a spatial pattern, strongly implying that *dpp* gene expression is regulated by regulators conveying spatial and temporal information in the MF (Fig 3E,E’, Fig S2D,D’). Third, high *dpp* gene expression appeared to depend on the activity of Ato. Indeed, we found that *ato* mutant cells located just anterior to wild-type Hh-expressing cells appeared to show reduced *dpp* transcription relative to wild-type cells (Fig S4). We therefore favor the view that the oscillatory expression of the *ptc* and *dpp* genes results from the pulsatile activities of Ato and/or Notch. This view remains to be tested.

### Reduced Hh signaling activity led to pattern irregularities along the MF

Our data so far indicated that *ptc* transcription and Ptc protein levels vary over time in phase with the Ato pulses and that Hh signaling activity positively regulates *ato* gene expression at the MF. Since Ptc negatively regulates Hh signaling, our observations raised the possibility that Hh might provide a diffusive temporal cue that coordinates *ato* gene expression along the MF. Here, we test whether changes in Hh activity impact the local synchrony of the Ato pulses along the MF. While synchrony would be best studied by live imaging, we found that Hh signaling perturbations appeared to affect the viability of cultured discs in our long-term imaging conditions. We therefore primarily studied local synchrony in fixed samples. Indeed, temporal differences in the pulsatile expression of Ato across columns, or in the periodic pattern of Notch activity, could in principle be detected in fixed samples in the form of irregularities in their expression pattern along the front. We first looked at the possible effect of increased Hh signaling activity in *mirr>ptc^RNAi^* eye discs and found no irregularities in the pattern of expression of the *ato* and *E(spl)* genes at the front (n=0/14; control discs, n=0/23; n, number of discs). In contrast, lowering Hh signaling in *mirr>hh^RNAi^* discs led not only to reduced *ptc* expression (Fig S5) but also to rare pattern irregularities (Fig 6A-B’; n=2/14; control discs, n=0/27). For instance, a lack of detectable *sca* expression at the position of an emerging IG (blue arrow, Fig 6B’; neighboring IGs indicated by yellow circles) suggested a temporal delay relative to its immediate neighbors, hence a defect in synchrony (*sca* was used in these experiments since the silencing of *hh* led to reduced *ato* expression, see Fig 1D-D’’). Likewise, the silencing of *ci* in *mirr>ci^RNAi^* discs resulted in occasional pattern irregularities along the front, with adjacent columns showing an abnormal phase difference (Fig 6B,B’; n=6/20). Finally, live imaging analysis of E(spl) dynamics using Halo-mδ, a knock-in version of E(spl)mδ (Couturier et al., 2026), showed pattern irregularities in *mirr>hh^RNAi^* discs indicative of a phase difference between adjacent columns (Figure 6C-C’’; note that Halo-mδ dynamics appeared to be normal in the control ventral compartment). Together, these occasional patterning defects that were indicative of local synchrony defects and that were observed in three different genotypes associated with low Hh activity supported the view that Hh contributes to coordinate fate dynamics along the MF.

**Figure 6.**
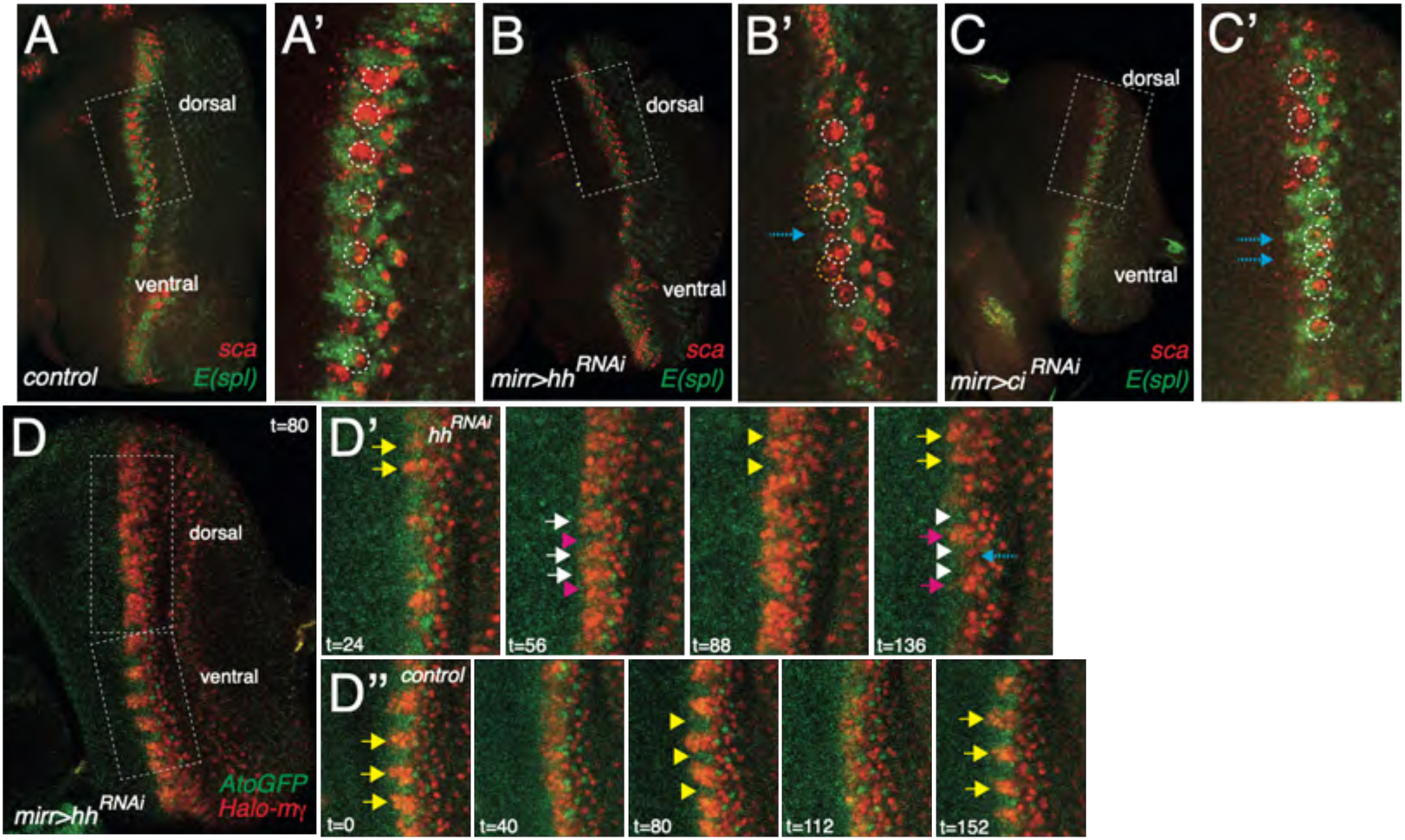
Pattern irregularities interpreted as coordination defects. A-B’) Analysis of fate dynamics in fixed eye discs stained for *sca* (red, exonic smiFISH probe) and *E(spl)* (green) in *mirr>+* (A,A’), *mirr>hh^RNAi^* (B,B’) and *mirr>ci^RNAi^* (C,C’) discs. High magnification views of the areas boxed in A-C are shown in A’-C’. In control discs (A,A’), Ato clusters (circles, A’) showed a continuous distribution of stages at the MF, from IGs (top) progressively resolving into R8s (bottom). In the *mirr>hh^RNAi^* disc shown in (B,B’), IGs expressing *sca* (yellow circles, B’) appeared to surround a soon-to-become IG that is not yet expressing *sca* (blue arrow, B’), indicative of delayed IG emergence. In the *mirr>ci^RNAi^* disc shown in (C,C’), two adjacent columns appeared to be in the same phase (a pair of blue arrows point to the two adjacent E(spl) teeth, each with a R8 cells at its base). D-D’’) Live imaging analysis of *mirr>hh^RNAi^* disc showing the dynamics of AtoGFP (green) and Halo-mγ (red) in snapshots (D’, D’’: high magnification views of the area boxed in D; time is in min). In control cells (D’’), ICs [E(spl) teeth, arrows] and IGs [Ato-positive clusters in between two E(spl) teeth, arrowheads] formed in synchrony (D’’). In *hh^RNAi^* cells (D’), most ICs and IGs appeared in synchrony (highlighted in yellow and white) but a delay was observed at one position: the teeth indicated by a blue arrow (t=136) showed a delay relative to neighboring teeth (in magenta).

## Discussion

Our study reveals that the pulsatile dynamics of Ato and Notch signaling along the front of differentiation in the developing eye of *Drosophila* is accompanied by the oscillatory expression of two Hh target genes, *ptc* and *dpp*. The latter oscillations did not appear to be generated by pulses of Hh synthesis since constant *hh* gene transcription was measured posterior to the MF in wild-type discs and since the Gal4-driven production of a GFP-tagged Hh by differentiated cells was sufficient to restore oscillatory *ptc* gene expression in *hh* mutant discs. Instead, these oscillations might be generated downstream of the pulsatile activities of Ato and/or Notch at the front of differentiation. This view is supported by the following observations: i) the transcription of the Hh target genes *ptc* and *rdx* in other area of the eye disc did not show oscillations; ii) *dpp* gene transcription in the MF appeared to be modulated by spatial and temporal cues present in the MF and iii) elevated *dpp* gene expression appeared to depend on the cell-autonomous activity of Ato; iv) finally, loss of Ptc did not prevent oscillatory *ptc* gene transcription. Together, these observations argue against the idea that oscillations might be generated via a molecular clock based on the negative regulation of Hh by Ptc with a time delay. Whether and how the pulsatile activities of Ato and/or Notch regulate the expression of the *dpp* and *ptc* genes remain, however, to be studied.

The periodic pulses of *ptc* transcription appeared to result in periodic changes in Ptc protein levels. Since Ptc interacts with extracellular Hh, these periodic variations in Ptc levels are predicted to result in the periodic depletion of extracellular Hh. Thus, Ptc may act in a non-cell autonomous manner by lowering extracellular Hh in a periodic manner. Since Hh contributes to regulate the activity of the *ato3’* and *ato5’* enhancer, this mechanism could contribute to synchronize the dynamics of *ato* gene expression across columns. Indeed, the diffusion of extracellular Hh across columns, the periodic changes in Ptc levels downstream of Ato and the binding of Hh to Ptc followed by its internalization should result in the periodic fluctuations of extracellular Hh. In turn, these temporal fluctuations may provide a temporal cue for the transcriptional regulation of the *ato* gene. Thus, Ato and Hh might form a pair of coupled periodic systems operating at the MF. The pulses of Ato would cell-autonomously regulate the periodic changes in Ptc levels while oscillatory Hh signaling would act in a cell non-autonomous manner to coordinate the column-autonomous dynamics of Ato across columns. In this model, the diffusion of Hh would provide a mechanism to couple the pulsatile expression of Ato across columns.

Beyond Hh, additional signals might contribute to coordinate cell behaviors locally. Previous studies have found that EGF receptor activity is high in IGs and, consistent with this, we observed that EGF receptor (EGFR) signaling showed pulses in the MF (Phan, Mestdagh *et al*., unpublished). Interestingly, these pulses of EGFR activity were Ato-dependent (Phan, Mestdagh *et al*., unpublished). However, whether pulsatile EGF signaling contributes to locally synchronize fate dynamics remains to be examined. Unfortunately, a direct role of EGF in regulating Ato is hard to test genetically because EGFR signaling is indirectly required to regulate the *ato* gene via its critical role in the expression of Hh (Rogers et al., 2005). Thus, whether EGFR acts in parallel to Hh to increase the robustness of fate patterning in the developing eye disc of *Drosophila* remains to be studied.

The traveling front of Hh in the developing eye offers similarities with the EGFR wave in the optic lobe of *Drosophila*. In this neuroepithelium, a proneural front self-propagates via a diffusive self-activating EGF signal. Activated EGFR triggers proneural gene expression and proneural factors down-regulates EGFR signaling by promoting a cell fate switch (Jörg et al., 2019; Sato and Yasugi, 2020). How fate transition is temporally coordinated along the front is not well understood but both the diffusion of EGF (Jörg et al., 2019; Sato and Yasugi, 2020) and cell-cell intercalations (Shard et al., 2020) likely contribute to smoothen this moving front of differentiation. Interestingly, oriented cell-cell intercalations were recently observed just anterior to the MF in the developing eye (Sui and Dahmann, 2024). Additionally, periodic cell flows were found to correlate with fate patterning in the eye (Gallagher et al., 2022).

Thus, tissue mechanics, possibly downstream of Hh (Corrigall et al., 2007), might also contribute to coordinate patterning dynamics along the front.

In conclusion, the regulation of a constant Hh signal by pulsatile Ptc is proposed to generate a temporal cue that is required to coordinate the dynamics of *ato* gene expression along the front of differentiation, thereby contributing to the precision of fate dynamics in the *Drosophila* eye.

## Acknowledgements

We thank H. Bellen, B. Hassan, D. Ish-Horowicz, Flybase, FlyORF, the Vienna Drosophila Ressource Center (VDRC) and the Bloomington Drosophila Stock Center (BDSC) for flies, plasmids, antibodies and/or database services. We thank L. Couturier and K. Mazouni (IP) for technical help. We are grateful to R. Levayer and all lab members for discussion and critical reading. We gratefully acknowledge UtechS Photonic BioImaging (Imagopole), C2RT, Institut Pasteur, supported by Agence Nationale pour la Recherche (ANR-10–INBS–04). This work was funded by grants from the Agence Nationale pour la Recherche to FS (ANR-10-LABX-0073 and ANR-22-CE13-0016-01).

## Methods

### Flies

The following transgenes and mutations were used: *ato^GFP^* (Mora et al., 2018), *ato3’-AtoGFP* (Couturier et al., 2026), *ato5’-AtoHalo* (Couturier et al., 2026), *E(spl)m*δ*^GFP^* (Couturier et al., 2026), *E(spl)m*γ*^GFP^* (Couturier et al., 2026), *E(spl)m*δ*^GFP^ E(spl)m*γ (Couturier et al., 2026), *hh^fse^* (BL-35562), *hh^bar3^* (BL-1376), *PBac[HhGFP]* (BL-86271), *UAS-HhGFP* (BL-81024), *UAS-hh^RNAi^* (BL-32489), *PBac[PtcCherry]* (BL-86272), *UAS-ptc^RNAi^* (BL-28795), *ci^RNAi^* (BL-28984), *mirr-Gal4* (*mirr^DE^*, BL-29650), *GMR-Gal4* (BL-1104), *FRT40A smo^3^* (BL-98355) and *FRT82B ato^1^*. Clones of *smo^3^* and *ato^1^* mutant cells were produced using the FLP-FRT system using *ey-FLP* (BL-5577). Adult flies were imaged using a Zeiss Discovery V20 stereo-macroscope using a 1.0X (PlanApo S FWD 60mm) objective.

### smiFISH and immunostaining

Fluorescent in situ hybridization was performed as described previously (Couturier et al., 2019). In brief, each smiFISH probe corresponds to duplexes between a set of gene-specific non-labelled primary oligonucleotides (probe set mix; equimolar mix of 23-29 different oligonucleotides with a 20 nt-long mRNA-binding moiety for a total length of 48 nt) which were annealed with a fluorescently labelled oligonucleotide (FLAP-X, 28 nt-long) coupled to Cy3, Alexa590 or Cy5. Exonic probes were used to detect the *ato, sca, E(spl)m*δ*/E(spl)m*γ [noted here *E(spl)*], *gfp* and *halo* mRNAs. Intronic probes were used to detect transcription foci at the *ptc*, *dpp*, *hh* and *rdx* loci. The oligonucleotide sequences of these probes are available upon request. To obtain smiFISH probes, FLAP-X oligonucleotides were annealed with the probe set mix in Tris-HCl 50mM pH=7.5, NaCl 100mM, MgCl2 10mM using a thermocycler (85°C, 3 min; 65°C, 3min; 25°C 3 min). All oligonucleotides were obtained from IDT Inc. Dissected tissues were fixed 20 min in 4% paraformaldehyde in PBS 1x, washed in PBS 1x and permeabilized in PBS 1x Triton X-100 0.5%. Discs were then washed twice in SSC 2x with Urea 4M before being incubated overnight (ovn) with the smiFISH probe at 37°C in SSC 2x, Urea 4M, Dextrane 10%, Vanydyl complex 10mM, 0.15 mg/ml salmon sperm DNA. Following this hybridization step, discs were sequentially washed in SSC 2x Urea 4M (2x at 37°c), then in SSC 2x (2x at room temperature) and in PBT 1x. For double smiFISH/immunostainings, primary antibodies were added in the hybridization mix and incubated ovn at 37°c. Secondary antibodies were incubated in PBT following all washes. For immunostainings, dissection of third instar larvae (in PBS 1x at room temperature), fixation (20 min in 4% paraformaldehyde in PBS 1x) and antibody staining in PBT (PBS 1X with Triton X-100 0.1%) were performed using standard procedures. The following antibodies were used: rat anti-ECad (mAb DCAD2, DSHB), mouse anti-Ptc (mAb Apa1, DSHB), guinea-pig anti-Sens (Nolo et al., 2000), mouse anti-Hry (a mix of the mAbs 24/1, 28/2 and 33/5) (Wainwright and and Ish-Horowicz, 1992), goat anti-GFP (ab6673, Abcam), rabbit anti-DsRed (#632496, Clonetech). Secondary antibodies were from Jackson’s laboratories. Stained discs were mounted in 4% N-propyl-galate, 80% glycerol. Images were acquired using a Zeiss LSM780 microscope with 40X (PL APO, NA 1.3, DIC M27) objectives and a Nikon Ti-E AX-NSPARC microscope with 40x (PL APO, NA 1.25).

### Live imaging

Live imaging was performed as described in (Couturier et al., 2026). Movies were acquired on a Nikon Ti2E spinning disk microscope equipped with a 40X (N.A. 1.15 Water WD 0.6) objective, a Yokogawa CSU-W1 spinning disk, a sCMOS Photometrics PRIM95B camera and 561/640 lasers. The AtoGFP and Halo-mγ signals were acquired every 10 mins in a tissue volume with a typical depth of 40 um (Dz=1.5um). We noticed that *mirr>hh^RNAi^* discs survived not as well as wild-type discs and could only be imaged for 3-4 hours at 25°C.

### Image analysis

To reconstruct pattern dynamics from fixed samples, each Ato cluster of the anterior-most row, row *n*, was staged and annotated as follows: IC/R8, 7; late IC, 0; interpulse, 1; early IG, 2; IG, 3; late IG, 4; interpulse, 5; early R8/IC, 6. Ato clusters at row *n-1* were automatically annotated by introducing a one period shift (see Fig S1). Each annotated cluster was then cropped along the direction perpendicular to the MF. The intensities of the *ato* and GFPmδ signals were scaled into [0,1] using min and max values measured within a ROI covering the entire MF. Cropped images were then scaled to the same size, grouped based on annotation and each group was averaged. The relative duration of each stage was estimated based on the number of clusters per annotated stage. This information was used to generate dynamic movie of 100 relative time points by smoothing over time the staged averaged images (sigma=5 time points). The dynamics of *dpp*, *ptc, rdx* and *hh* were reconstructed following the dynamics of *ato* and GFPmδ. Since the *dpp*, *ptc, rdx* and *hh* signals appeared as individual dots (produced by the hybridization of the intronic probes to the transcribed loci), these dots were identified by thresholding before averaging to reduce the background signal. In some cases, we observed that the low number of annotated clusters at stages 2 and 6 resulted in sampling bias during the averaging process. We therefore fused the clusters annotated as stages 2 and 3, and as stage 6 and 7 and proceeded as described above.

Kymographs were produced by computing the mean intensity of the signal along the x axis (AP) of the reconstructed dynamics. The resulting kymographs were smoothed by gaussian filtering (sigma=10 pixels).

To compare the reconstructed dynamics obtained from fixed samples with the one obtained by live imaging, we produced kymographs of the mean intensity of the signal along a line drawn centered on the E(spl) teeth or on the IG clusters as in (Couturier et al., 2026). The resulting kymographs were smoothed by gaussian filtering (sigma=5 pixels). The same approach was applied to study the spatial dynamics of *dpp* expression.

## Supplemental data

**Figure S1.**
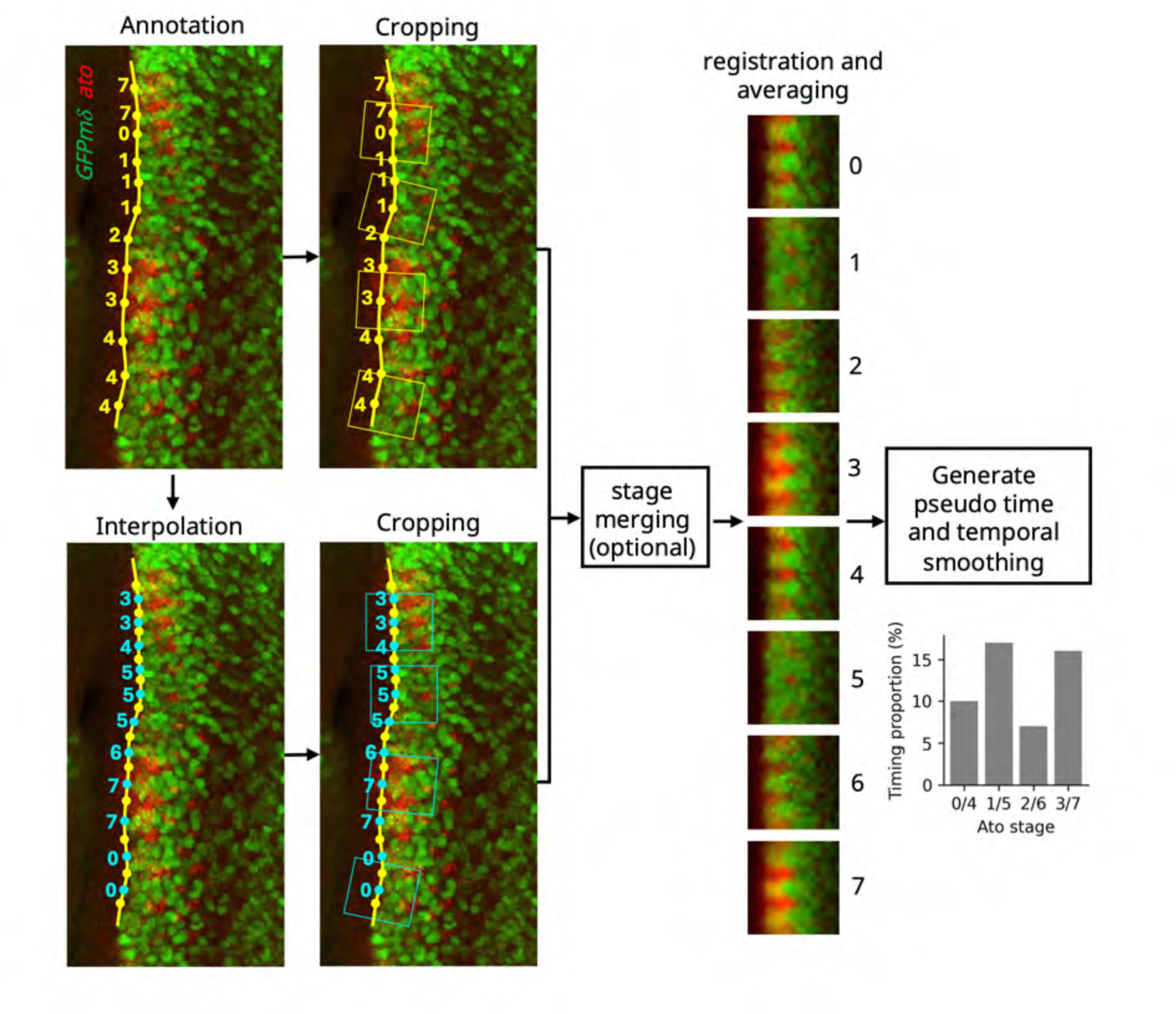
A pipeline for reconstructing gene expression dynamics at the MF from fixed samples. First (top left), starting from a maximal z-projected confocal image of an eye disc that was fixed and stained for *ato* mRNAs (red) and GFPmδ (green), we manually annotated the positions of the Ato clusters along the MF, and categorized these clusters into 8 stages, numbered 0 to 7 (see Methods for nomenclature). We then cropped the image, thereby producing images of staged clusters oriented perpendicular to the front (represented as yellow boxes; for better visualization, only one every third clusters are shown). Next (bottom left), we defined clusters corresponding to the previous row by interpolating positions and annotating categories automatically using a one period shift. Annotated clusters were then oriented and cropped as above (represented as cyan boxes). This pipeline produced a set of categorized images. Images of each category were then spatially registered and intensity-normalized before being averaged. For some datasets, categories corresponding to under-represented stages were merged to avoid sampling artefacts. This produced a set of 8 averaged images representing the dynamics. We next realigned these 8 frames along the AP axis to compensate for the progression of the MF, assigned a time duration for each of the 8 categories based on their relative frequency (see distribution plot, right) and smoothened the signals in time to reduce variability. This approach established the average spatial-temporal dynamics of Ato and E(spl)mδ at the MF (see movie 1).

**Figure S2.**
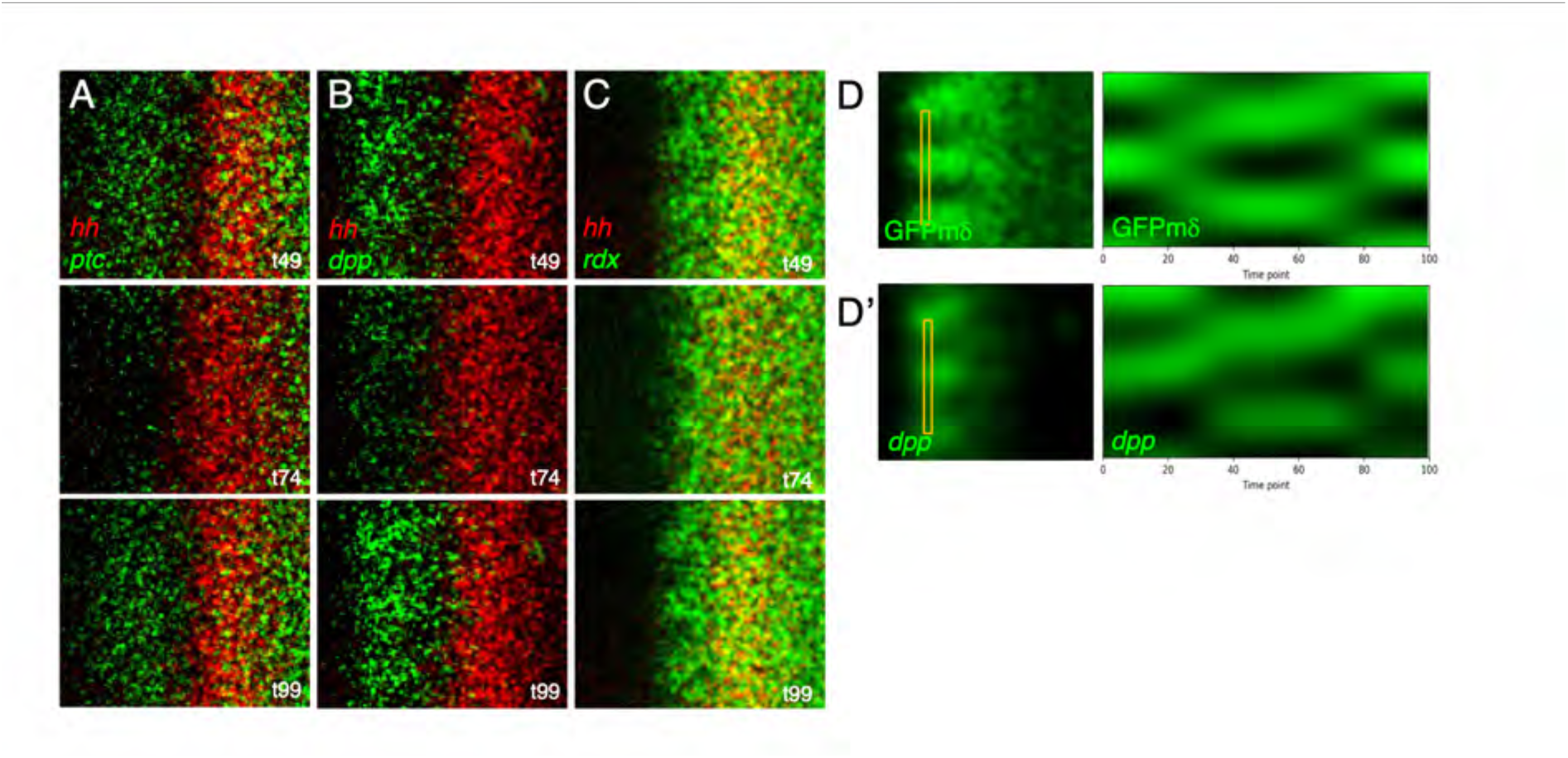
Transcription dynamics of the *hh*, *ptc*, *dpp* and *rdx* genes. A-C) Transcription dynamics of ptc (A, green), *dpp* (B, green) and *rdx* (C, green) relative to *hh* (red). For each gene, three snapshots taken from the reconstructed dynamics are shown. D,D’) Kymographs showing the spatial dynamics of GFPmδ (D) and *dpp* (D’). Kymographs were produced from the reconstructed dynamics using the area shown as yellow boxes (left, see Methods). This showed that the transcriptional pulses of *dpp* were spatially patterned.

**Figure S3.**
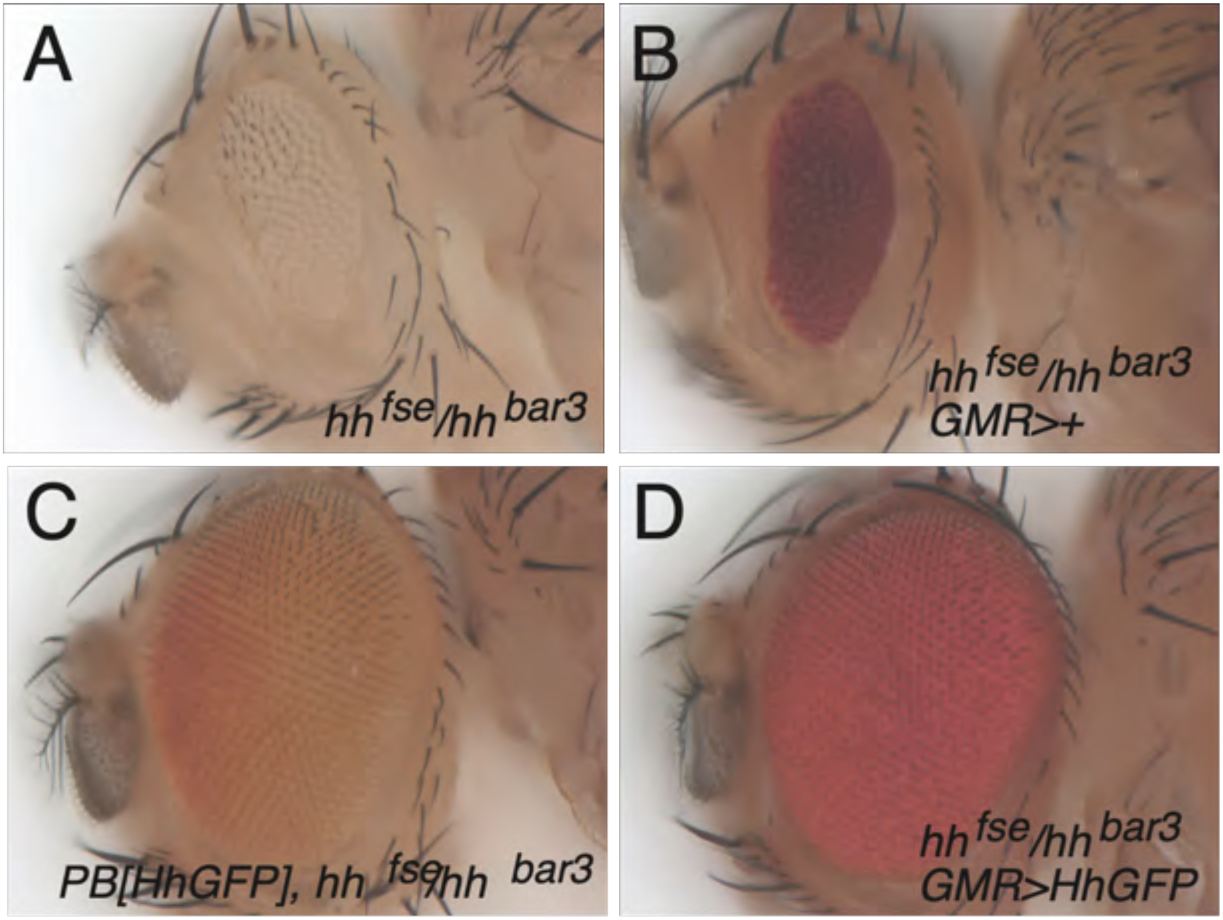
Rescue of the eye-specific *hh* loss function by HhGFP. A-D) The eye-specific loss of *hh* activity in *hh^fse^*/*hh^bar3^* flies led to a small eye phenotype in adult flies (A,B). This phenotype was rescued by a BAC-encoded version of GFP-tagged Hh (C) and by the expression of HhGFP in differentiated using the UAS/Gal4 system (D).

**Figure S4.**
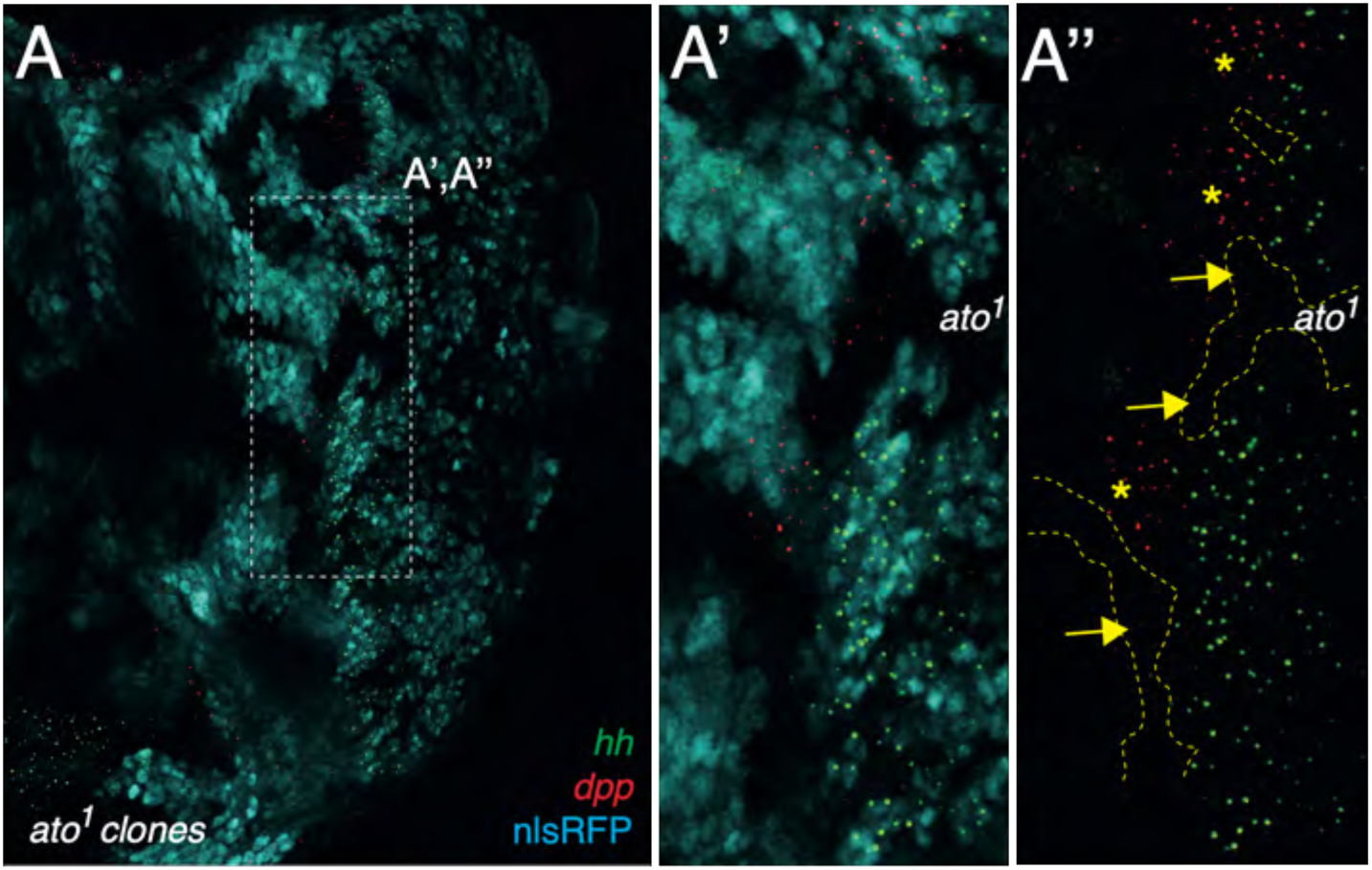
Ato-dependent transcription of *dpp* downstream of Hh. A-A’’) Analysis by smiFISH of *dpp* (red) and *hh* (green) transcription in eye imaginal discs carrying clones of *ato^1^* mutant cells marked by the loss of nlsRFP (cyan; high-magnification views of the region boxed in A are shown in A’A’’). MF cells that are mutant for *ato* (clone boudaries indicated by dotted lines in A’’) and that are located just anterior to wild-type cells expressing Hh showed reduced levels of nascent *dpp* transcripts (arrows in A’’) relative to wild-type MF cells (asterisks in A’’).

**Figure S5.**
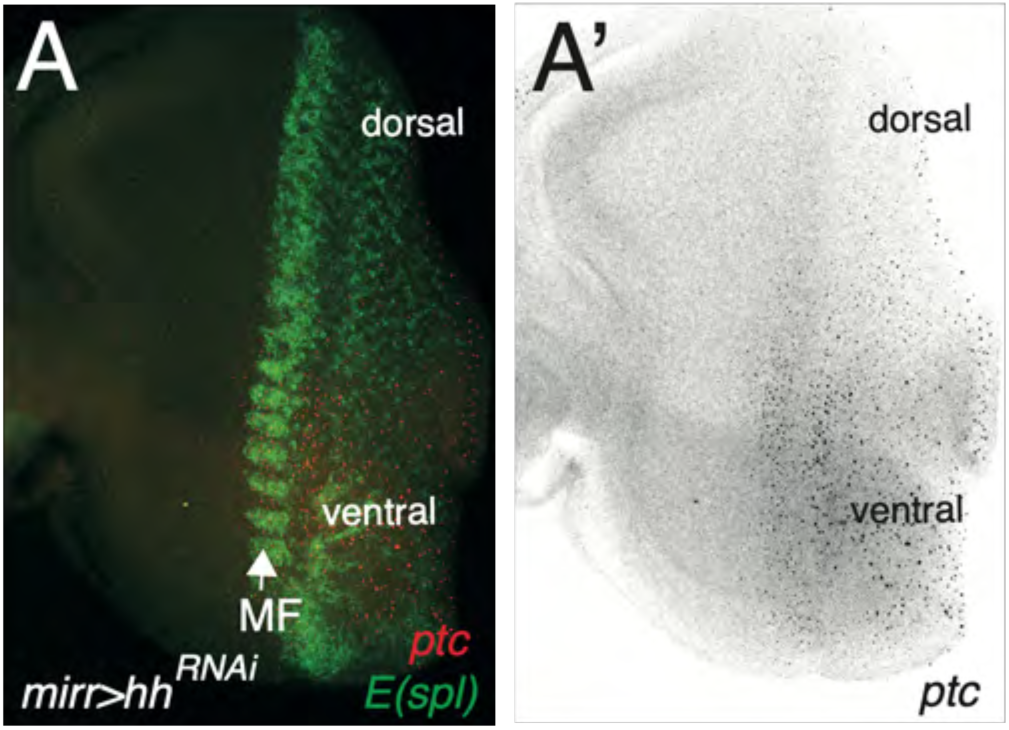
Low Hh signaling led to reduced *ptc* transcription. A,A’) The silencing of the *hh* gene in *mirr>hh^RNAi^* discs led to reduced *ptc* gene transcription (red in A; E(spl), green in A) in dorsal cells (MF position indicated by an arrow).

**Movie 1 – Reconstructed dynamics of Ato and E(spl)** – related to Fig 2 Reconstructed dynamics of *ato* mRNAs (red, smiFISH) and E(spl) (GFPmδ, green) over 2 consecutive pulses of Ato (time, arbitrary units)

**Movie 2 – Reconstructed dynamics of *ptc* transcription (with *ato*)** – related to Fig 3 Reconstructed dynamics of *ptc* transcription (green, smiFISH) and *ato* mRNAs (red, smiFISH) over 2 consecutive pulses of Ato (time, arbitrary units)

**Movie 3 – Reconstructed dynamics of *ptc* transcription (with E(spl))** – related to Fig 3 Reconstructed dynamics of *ptc* transcription (green, smiFISH) and E(spl) (GFPmδ, red) over 2 consecutive pulses of Ato (time, arbitrary units)

**Movie 4 – Reconstructed dynamics of *dpp* transcription (with ato)** – related to Fig 3 Reconstructed dynamics of *dpp* transcription (green, smiFISH) and *ato* mRNAs (red, smiFISH) over 2 consecutive pulses of Ato (time, arbitrary units)

**Movie 5 – Reconstructed dynamics of *dpp* transcription (with E(spl**)) – related to Fig 3 Reconstructed dynamics of *dpp* transcription (green, smiFISH) and E(spl) (GFPmδ, red) over 2 consecutive pulses of Ato (time, arbitrary units)

**Movie 6 – Reconstructed dynamics of *rdx* transcription (with ato)** – related to Fig 3 Reconstructed dynamics of *rdx* transcription (green, smiFISH) and *ato* mRNAs (red, smiFISH) over 2 consecutive pulses of Ato (time, arbitrary units)

**Movie 7 – Reconstructed dynamics of *hh* transcription (with ptc)** – related to Fig 4 Reconstructed dynamics of *hh* (red) and *ptc* (green) over 2 consecutive pulses of Ato (time, arbitrary units)

